# A persistent bacterial *Regiella* transinfection in the bird cherry-oat aphid *Rhopalosiphum padi* increasing host fitness and decreasing plant virus transmission

**DOI:** 10.1101/2024.09.26.615116

**Authors:** Wenjuan Yu, Qiong Yang, Alex Gill, Evatt Chirgwin, Xinyue Gu, Chinmayee Joglekar, Paul A. Umina, Ary A. Hoffmann

## Abstract

The bird cherry-oat aphid, *Rhopalosiphum padi*, is a major pest of agriculture due to its ability to directly damage crops and transmit plant viruses. As industries move away from chemical pest control, there is interest in exploring new options to suppress the impact of this pest. Here we describe the production of a transinfected line of *R. padi* carrying the bacterial endosymbiont, *Regiella insecticola*, originating from the green peach aphid, *Myzus persicae*. We show that *Regiella* increases the fitness of its novel host despite decreasing fitness in its native host. *Regiella* also shows a low level of horizontal transmission. Importantly the infection suppresses the ability of *R. padi* to transmit the barley yellow dwarf virus which damages wheat plants. This transinfection could be released to suppress virus transmission by aphids with its ability to persist and spread, making it potentially suitable for wide area release.

## Introduction

The bird-cherry oat aphid, *Rhopalosiphum padi*, is an important worldwide pest of several cereal crops, attacking wheat, barley, oat, rye, triticale and other cereals (Leather et al., 1989). The species has an invasive range that includes Europe, Africa, North America, South America, Australia and New Zealand, and normally shows cyclical parthenogenesis (Dixon, 1971) although in its invasive range and in the absence of its primary host it may only reproduce asexually (Valenzuela et al., 2010). *Rhopalosiphum padi* has a capacity to directly damage plants, but importantly can also vector many plant viruses, and in particular barley yellow dwarf virus (BYDV), which can be devastating to cereal crops. Control of this aphid is becoming increasingly challenging due to the evolution of chemical resistance (Gong et al., 2021; Tang et al., 2024; Wang et al., 2021) and the withdrawal of many pesticide registrations due to environmental concerns.

With increasing interest in developing non-chemical control options for this aphid and other pest species, renewed attention is focusing on utilizing and targeting bacterial endosymbionts that are common in aphids (Gao et al., 2023; Gu et al., 2023; Zindel et al., 2011). Most aphids are infected by the primary endosymbiont, *Buchnera aphidicola*, but a host of other endosymbionts are also found in aphids such as *Regiella insecticola, Hamiltonella defensa, Serratia symbiotica* and *Rickettsia* with other bacteria including *Arsenophonus, Rickettsiella viridis, Wolbachia* and *Fukatsia* being less common (Donner et al., 2024; Yang et al., 2023; Zytynska & Weisser, 2016). These bacteria can have a range of effects on their hosts, including the provision of nutrition (Zytynska et al., 2021), protection against parasitism and thermal extremes (Oliver et al., 2003; Rothacher et al., 2016), and fungal infections (Heyworth & Ferrari, 2015; Lukasik et al., 2013). Importantly endosymbionts can also be associated with both increased and decreased virus transmission (Higashi et al., 2023). However, the majority of these effects have been observed in studies investigating native endosymbionts of hosts, whereas proposed applications of endosymbionts in the control of other pest species have tended to focus on transinfections, which are generated from the introduction of an endosymbiont from one host species into another species (Gong et al., 2023; Gong et al., 2020).

Transinfections can result in quite different phenotypic effects, as typified by the strong reproductive and viral blocking effects induced by the *w*Mel *Wolbachia* infection in *Aedes aegypti* mosquitoes when sourced from *Drosophila melanogaster* flies (Walker et al., 2011).

In a recent overview of endosymbiont transfer experiments in aphids (Gu et al., 2024), several variables were identified that influenced the success of transinfections in aphids. The difficulties in achieving transinfections are mainly related to the fitness costs of new infections in host species, the absence of effective plant-based horizontal transmission, limitations of vertical transmission, and the presence of a high level of genetic variation in the endosymbiont, likely reflecting a past history of adaptation to a native host. Nevertheless, there has been a relatively high success rate of transinfections with some aphid endosymbionts, including *Rickettsiella* and *Regiella*. The former was transferred from its native pea aphid host *A. pisum* into *M. persicae*, resulting in substantial fitness costs and a change in aphid body colour (Gu et al., 2023). The latter has been successfully transferred from its native host *M. persicae* into *R. padi* as well as into Russian wheat aphid (*Diuraphis noxia*). The transinfection into *R. padi* is characterised here and is of particular interest due to the potential effects of *Regiella* on virus transmission in hosts (Higashi et al., 2023), coupled with the economic importance of *R. padi* as a vector of economically damaging plant viruses, which likely do more damage to cereal crops than direct feeding (Valenzuela & Hoffmann, 2015).

*Rhopalosiphum padi* was expected to be a suitable host for endosymbiont transfers because it harbours a wide range of secondary endosymbionts in China and Europe, including not only *Regiella* but also *Hamiltonella, Rickettsia, Spiroplasma* and *Arsenophonus*, and more rarely *Fukatsia, Serratia* and *Wolbachia* (Guo et al., 2019; Leybourne et al., 2023). In its invasive range in Australia and Chile, such secondary endosymbionts appear to be relatively rare (Yang et al., 2023; Zepeda-Paulo et al., 2018), providing an opportunity to introduce new endosymbionts into an invaded region of this pest. The phenotypic effects of native *Regiella* on *R. padi* have rarely been characterized (Thomsen et al., 2024) and this also applies to *Regiella* in other aphids. Some strains of *Regiella* may provide parasitoid protection, as shown for a native infection in *M. persicae* (Vorburger et al., 2010), for intraspecific transinfections across clones of the English grain aphid *Sitobion avenae* (Luo et al., 2020) and for interspecific transfers from *M. persicae* into pea aphid *Acyrthosiphon pisum* and the black bean aphid *Aphis fabae* (Jamin & Vorburger, 2019). However, other native *Regiella* strains in *M. persicae* (Vorburger et al., 2010) and in *A. pisum* do not provide protection (Oliver et al., 2003), while *Regiella* may increase predation risk from ladybirds in *R. padi* and *M. persicae* when tested at an mild temperature (Thomsen et al., 2024).

Beyond affecting the susceptibility of aphid hosts to natural enemies, *Regiella* may trigger host fitness costs and influence traits related to dispersal. In *Sitobion avenae*, the infection influenced the intrinsic rate of increase of cultures in a temperature-dependent manner, decreasing it at 25°C and 28°C but not necessarily at a higher temperature (Liu et al., 2019). In *M. persicae*, a native *Regiella* infection did not have strong fitness costs, while the same infection also did not induce costs when transinfected to two other aphid species (Jamin & Vorburger, 2019). *Regiella* may inhibit production of winged alate forms in *S. avenae* at 25°C which is expected affect movement rates of this aphid, though alate formation was not affected at higher temperatures (Liu et al., 2019). *Regiella* also inhibited alate formation in *A. pisum* when present as a native infection (Leonardo & Mondor, 2006) and a transinfection (Jamin & Vorburger, 2019).

Despite extensive screening of field populations, we have yet to detect *Regiella* in *R. padi* in Australia even though it is present in other aphids there (Yang et al., 2023). In this study, we developed an Australian line of *R. padi* transinfected with *Regiella* through microinjection of the native infection from *M. persicae*. We showed that the new *Regiella* (+) line of *R. padi* had a higher fitness as expressed by a shorter pre-reproductive period, increased reproductive output and extended longevity, in contrast to fitness costs or neutral effects on fitness associated with *Regiella* in other studies. We also found that this transinfection showed both vertical and horizontal transmission. Importantly, the infection suppressed BYDV acquisition and transmission. These impacts suggest that the transinfection could persist and may exert beneficial effects on disease control under some conditions.

## Methods

### Aphid cultures

A population of *R. padi* was provided by Dr. Piotr Trebicki (Grains Innovation Park, Horsham, Victoria). Populations were established from a single *R. padi* female and cultured on excised wheat seedlings (var. Trojan) 14 days after sowing (Zadoks GS12). These were inserted into cups (Fig. S1), which contained a water reservoir. Aphids were moved to new plants approximately every 2 weeks. All aphid cultures were kept in a controlled temperature (CT) room at 19 ± 1 °C with a 16:8 h light:dark photoperiod.

To increase aphid numbers for experiments, *R. padi* were taken from the cups and transferred to live wheat plants (GS20-25). The wheat plants and aphids were housed in insect rearing cages (30 × 30 × 62 cm, mesh 160 µm aperture, Australian Entomological Supplies Pty Ltd) for 3 weeks in a CT room at 19 ± 1 °C with a 16:8 h light:dark photoperiod. To age match individuals for experiments, adult apterous aphids were introduced to 60 mm Petri dishes with 1% agar and three excised wheat leaves (GS20-25, ~40 mm × 10 mm) embedded in 1% agar. Ten adults were introduced per Petri dish and placed in a CT room at 19 ± 1 °C with a 16:8 h light:dark photoperiod. After 24 h, adults were removed from the Petri dishes and nymphs allowed to develop for a specific time as required for each experiment.

### BYDV-PAV material

The BYDV-PAV serotype was provided by the Grains Innovative Park (Horsham, Victoria). To maintain the virus, wheat seedlings were sown in a CT room set at 20 ±1 °C, supported with growth lights (40W Grow Saber LED 6500K, 1200 mm length) and a 16:8 h light:dark photoperiod. After 18 days (GS13), ten aphids (mixed ages 3^rd^ instar to adult) were taken from the previous BYDV-PAV population, confirmed positive for BYDV, and placed into a plastic vial (15 mm diameter × 40 mm long) attached to a wooden skewer (100 mm long) and containing a second true leaf of wheat. The vial opening was plugged with cotton wool and the seedlings returned to the CT room. After 72 h, the aphids were removed and the seedling allowed to develop for a further 16 days. A leaf cutting (~30 mm × 10 mm) was taken to confirm virus infection through PCR (see Detection of BYDV). After confirmation of infection, plants were either used for experiments or aphids reintroduced to the viral wheat to repeat the process.

### Detection of Regiella

The detection of *Regiella* in aphids was assessed by quantitative PCR (qPCR) following total genomic DNA extraction using 5% Chelex 100 resin solution as described elsewhere (Yang et al., 2023). The qPCR was performed using a Roche LightCycler® 480 High Resolution Melting Master (HRMM) kit (Roche Diagnostics Australia Pty.Ltd., Castle Hill, Australia) and Immolase DNA polymerase (5 U/µL) (Bioline; Cat. No. BIO-21047) as described by Yang et al. (2023). Two primer sets were applied to amplify markers to confirm the quality of aphid DNA (actin as reference gene, actin_aphid_F1: GTGATGGTGTATCTCACACTGTC and actin_aphid_R1: AGCAGTGGTGGTGAAACTG) and the presence or absence of the *Regiella* infection (U99F: ATCGGGGAGTAGCTTGCTAC and 16SB4: CTAGAGATCGTCGCCTAGGTA). All samples were analysed concurrently with control samples, which had been previously confirmed as positive for *Regiella*. The presence of *Regiella* was assessed by comparing the melting temperature (Tm) of each sample with positive controls. Crossing point (Cp) values of two*–*three consistent replicate runs were averaged. Differences in Cp values between the actin and the target endosymbiont markers were transformed by 2 ^-ΔΔ Cp^ to produce relative density measures.

### Detection of BYDV

The first step in BYDV detection was to extract the total RNA from individual aphid and plant samples using a Monarch Total RNA Miniprep Kit (NEB, Ipswich, MA, USA) as described elsewhere (Chirgwin et al., 2023). RNA from each sample was then reverse transcribed into cDNA using a high-capacity cDNA reverse transcription kit (Thermo Fisher, Waltham, MA, USA), which was then used as the template for qPCR assays as described above (Detection of *Regiella*). Two primer sets were used to amplify markers specific to BYDV-PAV (BYL: GTGAATGAATTCAGTAGGCCGT; BYR: GTTCCGGTGTTGAGGAGTCT) and wheat β-actin (actin_wheat_F1: CGGTATCGTAAGCAACTG; actin_wheat_R1: GCGTATCCTTCGTAAATGG) (Paolacci et al., 2009). Data was analysed using 2 ^-ΔΔ Cp^ method as noted above. A semi-quantitative PCR assay using conventional PCR was performed when the overall virus density was lower than the detection limit of the qPCR assay using the same program and primers with qPCR. Each band from the 2% agarose gel was given a score based on a 1-6 scale from a set of standard images, with 1 being the lowest and 6 being the highest.

### Endosymbiont transfer through microinjection

*Regiella insecticola* (Enterobacterales: Enterobacteriaceae; henceforth referred to by genus) from a naturally infected *M. persicae* was collected from capsicum in Melbourne, Australia (–37.830290, 145.023525) and introduced to adult *R. padi* through microinjection using the method outlined in Gu et al. (2023). After microinjection, aphids were reared on 35 mm Petri dishes with 1% agar and excised wheat leaves (~30 mm × 10 mm). After producing more than four nymphs, the adult females were removed, stored in 100% ethanol and screened for *Regiella* infection. Nymphs with infected parents were kept and this process repeated until a 100% infection rate was achieved in the third generation. Once the *Regiella* infection was stable, the aphids were maintained following the methods described above. The *Regiella* positive line (+) was screened by qPCR every month and prior to commencing each experiment. A colony of aphids uninfected by *Regiella* (−) but derived from the same original clone was maintained alongside the transinfected line.

### *Horizontal transmission of* Regiella

To assess horizontal transmission of the infection between *Regiella* (+) and *Regiella* (−) aphids, two treatments were considered: (1) low density exposure on a Petri dish (Fig. 1A), and (2) high density using living plants (Fig. 1B). We ensured all aphids tested in our study were the same age and life stage by setting up ten age-matching plates for each of the *Regiella* (+) and *Regiella* (−) lines. Each age-matching plate consisted of 30 adult aphids within a 100 mm petri dish containing *T. aestivum* leaves placed in 10 g/L agar. The first treatment involved the mixing of a two-day-old *Regiella* (+) nymph and a two-day-old *Regiella* (−) nymph in a Petri dish with 1% agar and an excised wheat leaf (GS25; ~20 mm × 10 mm). Fifty replicate Petri dishes were established in a CT room at 19 ± 1 °C with a 16:8 h light:dark photoperiod.

**Figure 1.**
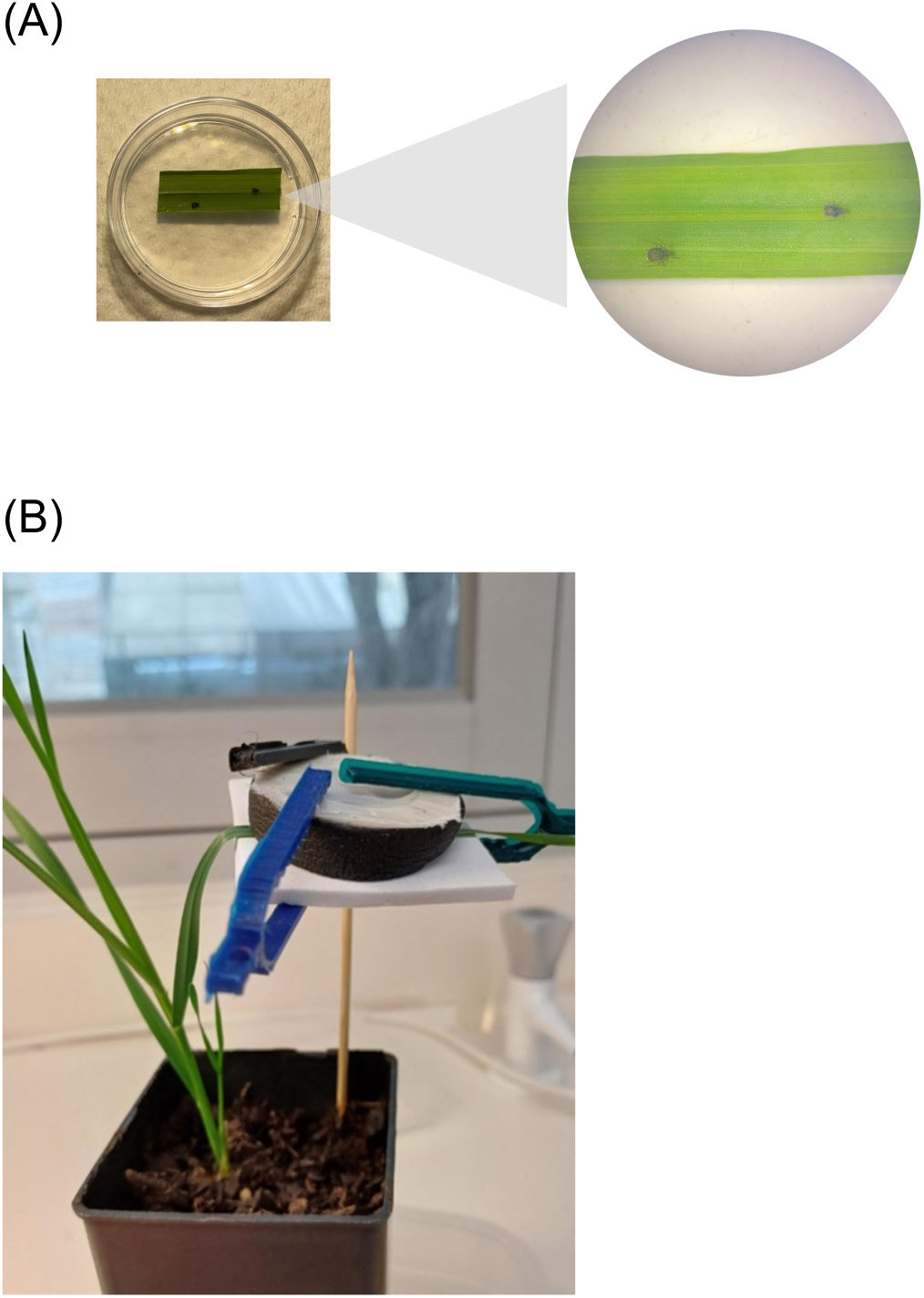
Horizontal transmission experiments at low and high density. (A) Set up for low density horizontal transfer experiment. A single *Regiella* (+) aphid was mixed with a *Regiella* (−) aphid in a Petri dish containing an excised wheat leaf. (B) Set up for high density horizontal transfer experiment. 20 *Regiella* (+) and *Regiella* (−) aphids were mixed and held via a clip cage attached to a living wheat plant.

Aphids were left to develop for 10 days before both individuals were collected and later screened for *Regiella* infection. At the point of reproduction, aphids were checked every 24 h and all nymphs removed. In the second high density treatment, 20 *Regiella* (+) and 20 *Regiella* (−) two-day-old nymphs were mixed in a clip cage (40 mm diameter × 1 mm high and a 25 mm internal diameter) attached to the second-true leaf of a living wheat plant (GS13) in the same maintenance condition with the first treatment. Aphids were held together for 72 h before being removed and screened for *Regiella* infection. This process was repeated twice.

### *Effect of* Regiella *infection on life history of apterous and alate* R. padi *across multiple generations*

Three *Regiella* (+) one-day-old nymphs and 3 *Regiella* (−) one-day-old nymphs were transferred into separate Petri dishes (35 mm × 10 mm), each containing a layer of 1% agar and 3 excised wheat leaves (~20 mm × 10 mm). After developing into apterous adults, aphids were checked every day and allowed to produce nymphs (G1). Forty-five one-day-old G1 individuals were then used to measure history life traits. These nymphs were placed individually into Petri dishes with a single wheat leaf (~20 mm × 10 mm) on 1% agar. All Petri dishes were placed in a CT room at 19 ± 1 °C with a 16:8 h light:dark photoperiod. Aphids were transferred to new Petri dishes twice a week and aphid survival assessed daily, for 7 days. Once aphids reached adulthood, they were separated into alate and apterous adults and left to produce nymphs (G2). These nymphs were counted and removed from the Petri dishes daily to determine the time to reproductive maturation time, longevity, lifetime fecundity and duration of reproductive period of the G1 individuals.

We ran this experiment twice, measuring the above traits at the 4^th^ generation post transinfection and the 10^th^ generation post transinfection. These experiments are not directly comparable due to some differences in equipment and the conditions under which they were undertaken (nature of incubator, plants) but the aim was to test the consistency of patterns between the *Regiella* (+) and *Regiella* (−) lines.

### *Impact of* Regiella *on barley yellow dwarf virus acquisition and transmission*

We ran two experiments to explore acquisition and transmission efficiency between the *Regiella* (+) and *Regiella* (−) aphids. In the first experiment, we tested how *Regiella* altered the density of BYDV acquired by *R. padi* after feeding on infected plants. In the second experiment, we measured both acquisition and transmission efficiencies (Fig. 2).

**Figure 2.**
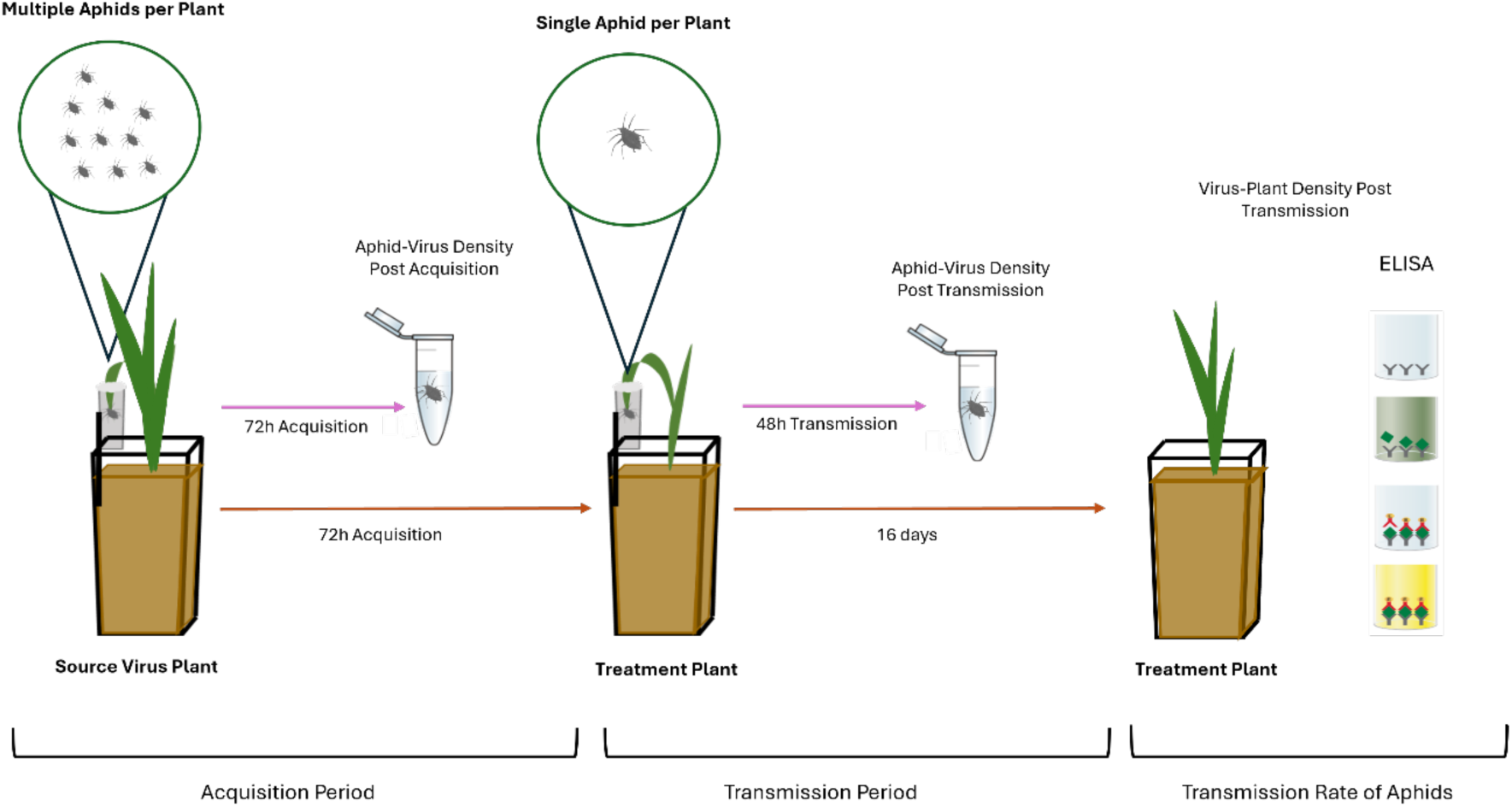
Virus transmission experiment to examine the ability of *R. padi* to acquire and transmit BYDV-PAV.

To establish sufficient viral source material for the first experiment, wheat plants were inoculated using the methods outlined earlier in the ‘BYDV-PAV material’ section. *Regiella* (+) *and Regiella* (−) aphids were then age matched on Petri dishes as described above (Horizontal transmission of *Regiella*). The resulting two-day-old nymphs were introduced directly to BYDV-infected wheat or BYDV-uninfected wheat plants in pots. Ten aphids were placed into a plastic vial attached to a wooden skewer and containing a second true leaf of wheat (see “Acquisition period” of Fig. 2, replicated 10 times per aphid line). The aphids were left for 72 h to acquire the virus. The experiment was conducted in a CT room at 19 ± 1 °C with a 16:8 h light:dark photoperiod. Aphids were then removed and stored in RNAlater^Tm^ solution (Invitrogen) for qPCR screening of aphid virus density post-acquisition.

In the second experiment, we age-matched aphids as described above and tested the density of BYDV in *R. padi* following acquisition. Here, we placed ten non-viruliferous aphids within a clip cage (Fig. 1B, rather than directly onto a plant) and allowed them to feed on BYDV-infected wheat plants for 48 h in a CT room at 19 ± 1 °C with a 16:8 h light:dark photoperiod. We established 15 replicate plants with *Regiella* (+) aphids and 15 replicate plants with *Regiella* (−) aphids and allowed them to feed for 48 h before removal, storage in RNAlater^Tm^ and later screening.

Next, we tested the density of BYDV in wheat plants following inoculation by viruliferous *R. padi*. We first inoculated *Regiella* (+) and *Regiella* (−) aphids with BYDV by allowing them to feed on infected wheat plants for 4 days. Aphids were inoculated in groups of 10 per plant. Each group of 10 viruliferous aphids was then placed in a clip cage and allowed to feed on uninfected wheat plants for 48 h before removal, storage in RNAlater^Tm^ and later screening. The plants were grown for a further 14 days to allow the virus to replicate, after which time the entire plant was removed for BYDV quantification using a DAS (double antibody sandwich)-ELISA kit from BIOREBA AG (Reinach, Switzerland), following the manufacturer’s instructions. The ELISA assays were analyzed on a Byonoy absorbance microplate reader (Byonoy GmbH, Germany) at 405 nm with positive readings at least three times the negative control. We established 24 replicate plants for transmission by *Regiella* (−) aphids and 23 replicate plants for *Regiella* (+) aphids, and obtained virus density readings in 15 plants exposed to *Regiella* (−) aphids and 9 plants exposed to the *Regiella* (+) aphids.

### Statistics

Where percentages were computed (e.g. rates of horizontal transmission), 95% confidence intervals (95% CIs) were obtained through Epitools (https://epitools.ausvet.com.au/) using the Wilson method. During the horizontal transmission experiment, one or both aphids died in 11 (out of 50) Petri dishes, and so these were excluded from the analysis. For the life history traits including time to maturation, we compared the data for the alate and apterous aphids separately using *t*-tests, except in the case of the duration of the reproductive period, where we used non-parametric Mann-Whitney tests because the data were skewed. All tests were run using IBM Statistics SPSS version 24. For the virus acquisition and transmission experiments, we compared the *Regiella* (+) and *Regiella* (−) treatments using generalized/general linear models run in the statistics program *R*.

We also used the time to reproductive maturation (refer to here as the pre-reproductive period) and fecundity to estimate *r*^*m*^, the intrinsic rate of increase of a population, following the procedure suggested for aphids (Wyatt & White, 1977). This method uses the time to maturation, *d*, and the number of progeny produced in a period equivalent to *d*, defined as *M^d^*, in the equation *r^m^* = 0.74(log*^e^M^d^*)/*d*. Bootstrapping of the original data was used to produce confidence intervals based on 2,000 bootstrap values.

## Results

### *Horizontal transmission of* Regiella

In the qPCR assay screening for *Regiella*, aphids with Tm values of *Regiella* amplification within the range of positive controls (86.63-87.33) were considered *Regiella* positive. Aphids with *Regiella* relative (compared to the actin gene) density > 0.1 were considered to carry a high density of bacteria, while aphids with a *Regiella* density < 0.1 were considered to carry a low density of bacteria. Aphids were considered to lack the bacterium if Tm values of *Regiella* amplification were not within the range of positive controls.

We found horizontal transmission occurred between aphids on wheat leaves within a Petri dish (in the low-density treatment), although at a low rate of 15% (95% CIs: 7.25%, 29.73%). The recipient uninfected aphids testing positive for *Regiella* had a low average *Regiella* density based on the above criteria, with an estimated relative density of 0.02 (95% CIs: 0.001, 0.06) in contrast to the positive aphids which had an average density of 25.00 (95% CIs: 3.04, 42.08).

For the high-density treatment in clip cages on living plants, we screened 87 aphids, of which 44 had a relative density > 0.1 indicating that these had a high *Regiella* density (and likely include the original *Regiella* (+) aphids). Of the remaining 43 individuals, 38 were found to be infected by *Regiella*, which leads to an estimated horizontal transmission rate of 88.37% (95% CIs: 75.52%, 95.93%). Average density of *Regiella* based on the 2 ^-ΔΔ Cp^ method was 0.02 (95% CIs: 0.0004, 0.0627) for putative *Regiella* positive aphids that acquired *Regiella* via horizontal transmission and 109.81 (95% CIs: 7.78, 5293.48) for the remaining aphids with high *Regiella* density.

### Regiella infection improves fitness components in R. padi

We measured life history traits in *Regiella* (+) and *Regiella* (−) *R. padi* on two occasions, at the 4^th^ generation and the 10^th^ generation post transinfection. At the 4^th^ generation, we observed a sharp increase in fitness in both *Regiella* (+) apterous and alate aphids compared with *Regiella* (−) aphids, including for pre-reproductive period (t=7.943, df=63, *P* < 0.001; t=9.788, df=57, *P* < 0.001, Fig. 3A), longevity (t=17.3, df=63, *P* < 0.001; t=13, df=57, *P* < 0.001, Fig. 3B) and fecundity (t=15.8, df=63, *P* < 0.001; t=17.62, df=57, *P* < 0.001, Fig. 3C). The differences in fecundity reflected a longer reproductive duration ((t=5.609, df=63, *P* < 0.001; t=16.51, df=57, *P* < 0.001, Fig. 3D) and a higher peak fecundity as well as a shorter period to reach reproductive maturation in *Regiella* (+) aphids. We estimated *r*^*m*^ from the pre-reproductive period and reproductive output over the same period; for the alates, the *r*^*m*^ for *Regiella* (−) aphids was 0.149 (95% CIs: 0.134, 0.161) compared with a *r*^*m*^ of 0.273 (95% CIs: 0.268, 0.290) for *Regiella* (+) aphids. For the apterous aphids the *r*^*m*^ values for *Regiella* (−) aphids was 0.275 (95% CIs: 0.261, 0.286) compared with a *r*^*m*^ value of 0.376 (95% CIs: 0.363, 0.388) for *Regiella* (+) aphids. These values highlight an overall advantage of the *Regiella* (+) aphids with non-overlapping confidence intervals.

**Figure 3.**
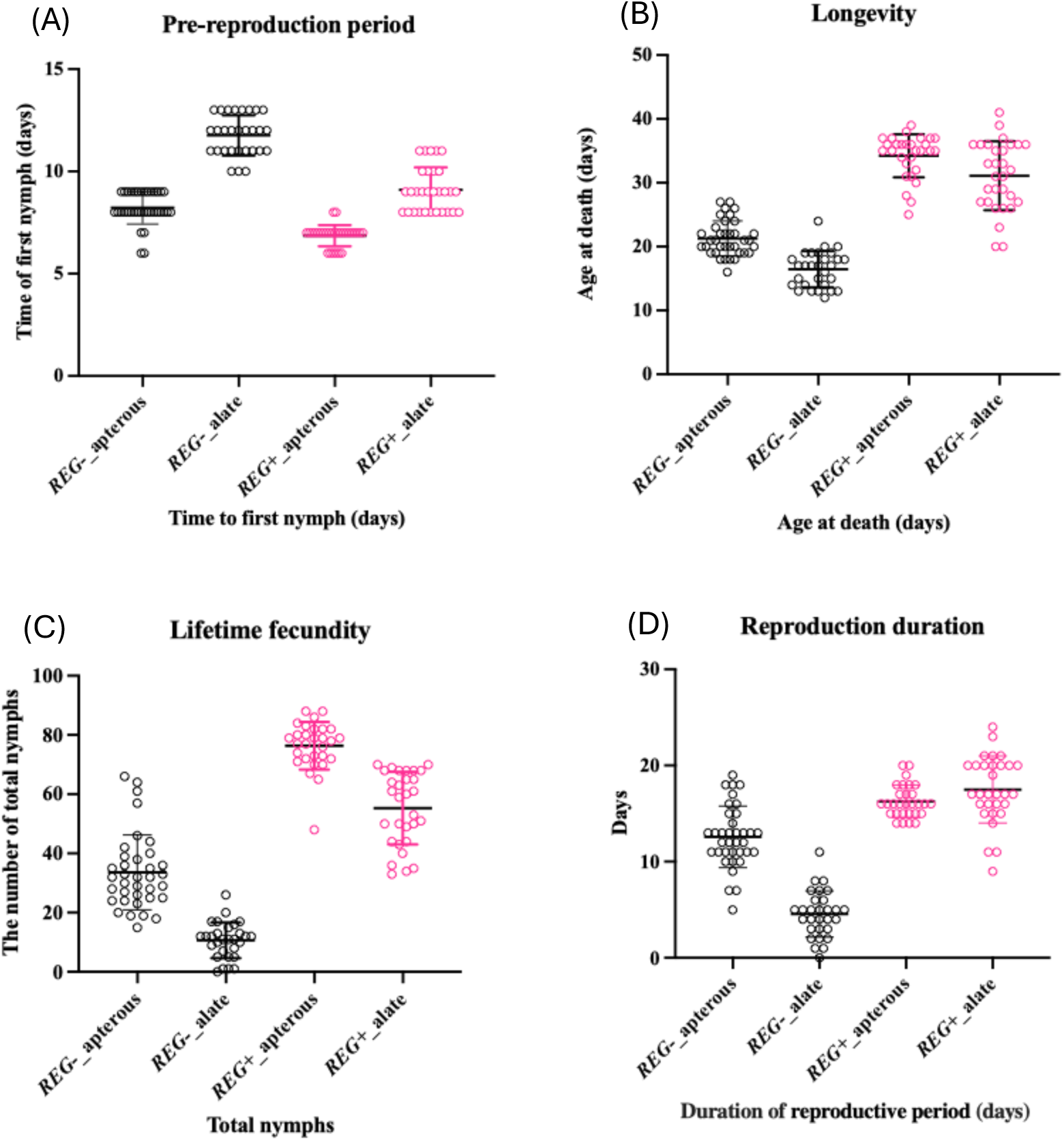
Life history traits of *Regiella* (+) and *Regiella* (−) aphids at the 4th generation post transinfection. Aphids were assessed for (A) pre-reproductive period, (B) longevity, (C) lifetime fecundity and (D) duration of reproductive period. Aphids were separated into alate and apterous groups.

We repeated these assessments at the 10^th^ generation post transinfection with mostly similar results (Fig. 4). *Regiella* (+) apterous aphids had a shorter time to reproductive maturation (t=3.95, df=68, *P* < 0.001, Fig. 4A) and higher fecundity (t=4.409, df=68, *P* < 0.001, Fig. 4C). Both apterous and alate aphids had an increase in longevity (t=5.212, df=68, *P* < 0.001; t=3.359, df=53, *P* < 0.05, Fig. 4B), and a longer reproductive duration (t=3.115, df=68, *P* < 0.05; t=2.777, df=53, *P* < 0.05, Fig. 4D) compared with *Regiella* (−) aphids. However, for alates there was no difference in the pre-reproductive period or lifetime fecundity. For alates, the *r*^*m*^ value for *Regiella* (−) aphids was 0.146 (95% CI 0.136, 0.161) compared with a *r*^*m*^ of 0.266 (95% CI 0.253, 0.268) for *Regiella* (+) aphids. For apterous aphids, the *r*^*m*^ value for *Regiella* (−) aphids was 0.272 (95% CI 0.262, 0.309) compared with a *r*^*m*^ of 0.356 (95% CI 0.351, 0.373) for *Regiella* (+) aphids.

**Figure 4.**
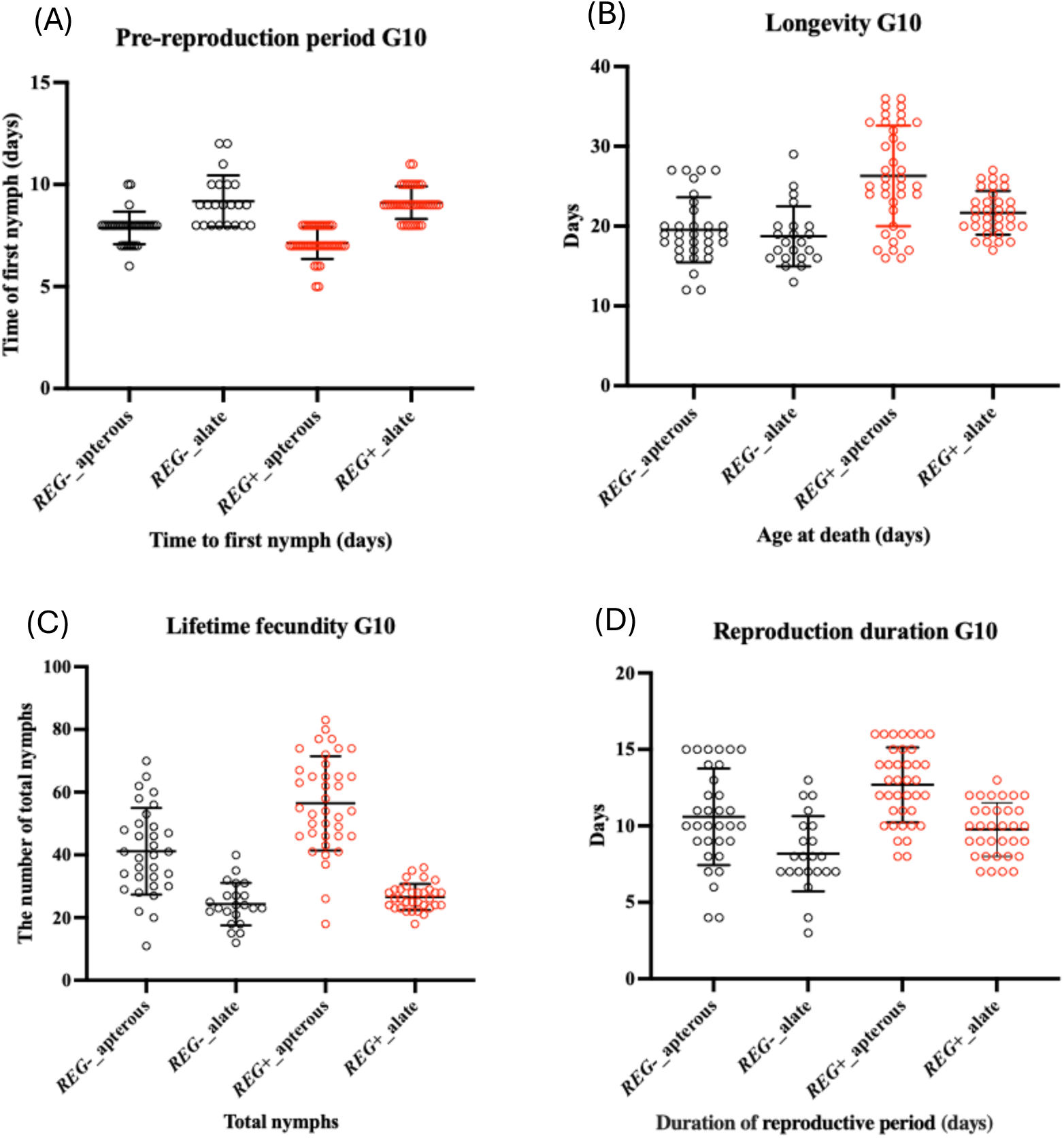
Life history traits of *Regiella* (+) and *Regiella* (−) aphids at the 10th generation post transinfection. Aphids were assessed for (A) pre-reproductive period, (B) longevity, (C) lifetime fecundity and (D) duration of reproductive period. Aphids were separated into alate and apterous groups.

### *Impact of* Regiella *on transmission and acquisition of BYDV*

We screened for BYDV in 144 aphids (72 *Regiella* (+) and 72 *Regiella* (−) individuals), which were exposed to the virus-infected source plants to acquire virus. While the actin reference gene amplified as we observed previously, viral density was under the detection limit of the qPCR assay. We then screened the aphids using conventional PCR. There were only 5 out of 77 *Regiella* (+) aphids clearly infected by BYDV, with very low virus density as judged optically, while 32 out of 77 *Regiella* (−) aphids were infected (contingency test, χ^2^ = 28.71, d.f. = 1, *P* < 0.001), with a much higher density (generalized linear model, χ^2^ = 22.989, d.f. = 1, *P* < 0.001) compared with that in *Regiella* (+) aphids. The optical density data averaged across three assessments is provided in Fig. 5. These screening result suggests that the *Regiella* infection reduces BYDV acquisition in *R. padi* when virus density in the plants is low.

**Figure 5.**
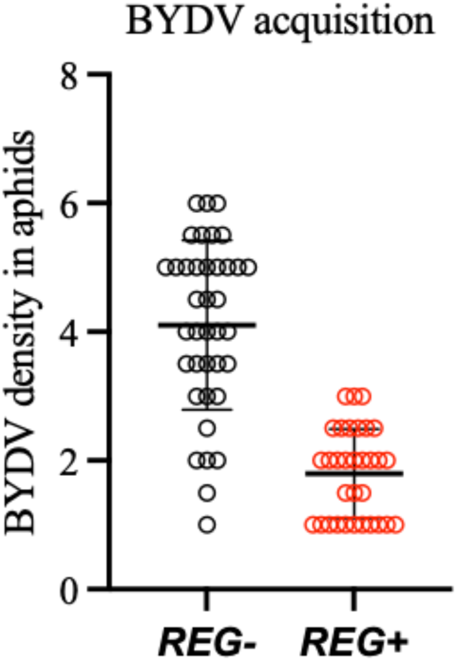
BYDV density in *Regiella* (+) and *Regiella* (−). Density data were obtained by optical scanning of bands from conventional PCR.

In the second experiment, we again observed a lower rate of BYDV acquisition in *Regiella* (+) aphids (Fig. 6a; χ^2^= 8.132, d. f.= 1, *P* = 0.003). The difference between strains was smaller than in the first experiment, but it is worth noting that here we are scoring density via qPCR which we have found to produce more variable results than conventional PCR. We also found significantly higher BYDV optical density in the 15 out of 24 wheat plants exposed to *Regiella* (−) aphids which became viruliferous and 9 plants out of 23 plants exposed to *Regiella* (+) aphids which became viruliferous (Fig 6b; F^1,22^ = 1.4043, *P* = 0.042 when non-viruliferous plants excluded), suggesting that transmission has also been affected.

**Figure 6.**
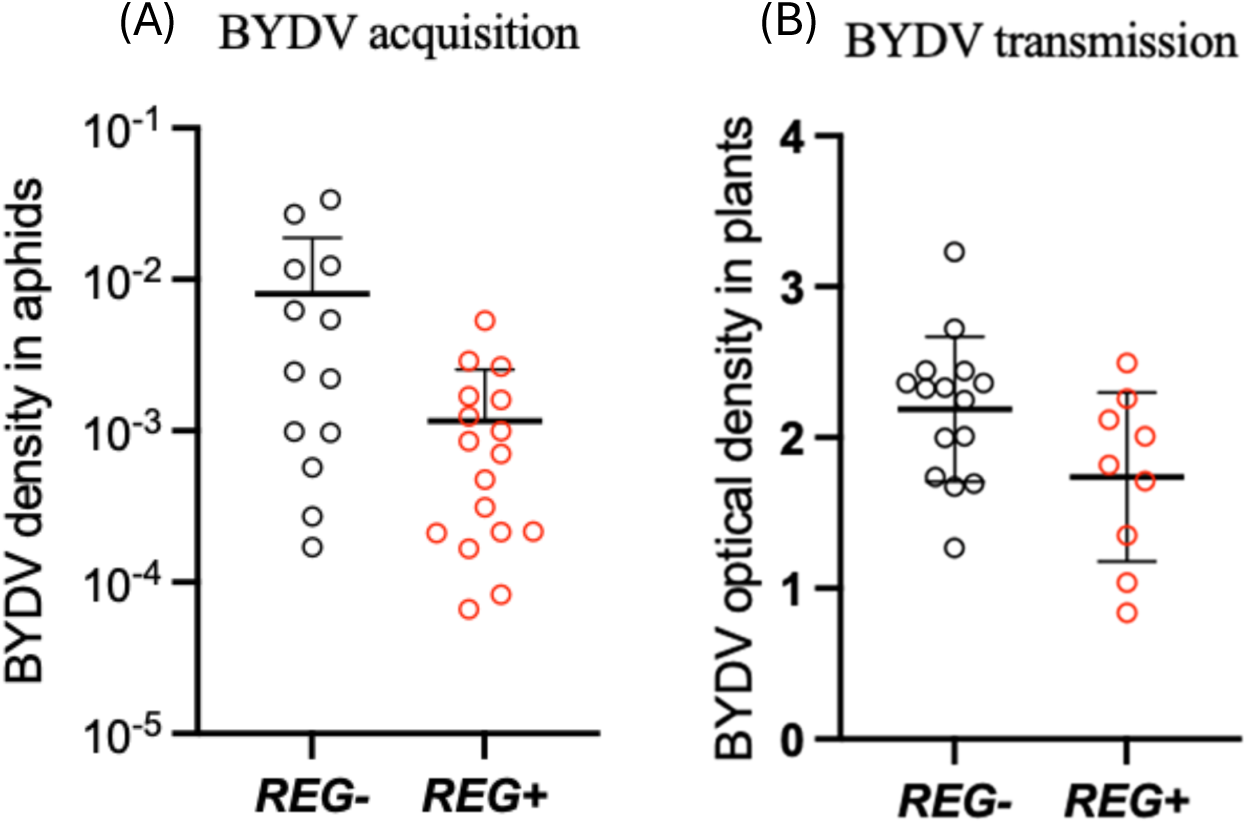
Acquisition and transmission of BYDV by *R. padi* when virus is at a high level in wheat plants. (A) Acquisition of virus as measured by qPCR. (B) BYDV density in plants as measured by ELISA where transmission to wheat plants was successful. Aphids carried the *Regiella* infection (REG+) or were uninfected (REG−).

## Discussion

Here we show that the endosymbiont infection *Regiella* can be successfully transinfected from its native *M. persicae* to a different aphid. Transinfections with this endosymbiont have been achieved before, with 4 successful documented interspecific transfers (e. g. Łukasik et al., 2015; Tsuchida et al., 2011) but there have also been failures such as the failure to transfer *Regiella* from *Macrosiphum euphorbiae* to other aphids (Gu et al., 2024). *Regiella* can also be lost following a change in host plant (such as the loss of *Regiella* when pea aphids were shifted from subterranean clover to *Medicago sativa* and through poor vertical transmission as in *Regiella* from pea aphids when transferred to *M. persicae* and *R. padi (*Gu et al., 2024). These findings highlight the importance of donor strain in successful transinfections with *Regiella* with success being affected by factors such as stability of the endosymbiont in the donor strain and relatedness of the donor and target host species (Gu et al., 2024; Jamin & Vorburger, 2019; Łukasik et al., 2015).

Although we have evidence of horizontal transmission of *Regiella* particularly when infected and uninfected aphids are held together at a high density on the same leaf, we are uncertain about whether there is “true” plant-based transmission which would require physical separation of the infected and uninfected aphids and direct evidence of *Regiella* being present in leaf tissue. In the case of *Rickettsiella* transinfected into *M. persicae* and tested on bok choy, we ran several experiments to show that transmission occurred when infected and uninfected aphids were physically separated even when leaves were on different stems (Gu et al., 2023). In our current work, transmission may occur through contact as well as plant tissue. However, plant-based transmission is known for other aphid endosymbionts and particularly *Hamiltonella* that can be horizontally transmitted in several cereal aphids including *R. padi* (Li et al., 2018). In *Sitobion miscanthi* feeding on wheat, *Hamiltonella* persisted locally on plant tissue for several days and, following transfer, lines showed stable vertical transmission (Li et al., 2018), something we also observed in lines of *R. padi* infected by *Rickettsiella* (Gu et al., 2024).

Newly acquired facultative endosymbionts are often seen as parasites, gaining resources from their hosts to increase their own fitness (Hoffmann & Cooper, 2024). However, the current findings also highlight that fitness effects of an endosymbiont in a new background can be unpredictable. Here the same endosymbiont infection (*Regiella*) from M. persicae has positive effects when transinfected to a new aphid host. In its native host, we have previously found that this infection does not generate any obvious fitness costs (Yang and Gill, unpublished) which is also consistent with observations for another native *Regiella* strain in *M. persicae* originating from Australia (Jamin & Vorburger, 2019). The positive effects of *Regiella* on host fitness may help it to initially spread and persist within host populations, although endosymbiont–host interactions and dynamics in the field are expected to be complex, typically depending on environmental conditions and genetic background of the host (Hoffmann & Cooper, 2024).

Importantly our data point to the inhibition of plant virus transmission associated with the *Regiella* transinfection in *R. padi*. Previous work has shown impacts of endosymbionts on host viruses, such as native *Regiella* reducing the impact of APV virus that was harmful to its pea aphid host (Higashi et al., 2023). Previous research has also demonstrated densovirus inhibition in the wheat aphid *Sitobion miscanthi* associated with the native symbiont SMLS which induced antiviral factors in the aphid host to prevent the expression of negative fitness effects associated with the virus (Li et al., 2024). Inhibition of the transmission of plant viruses has also been demonstrated in brown plant hoppers transinfected with *Wolbachia* from a related species (Gong et al., 2020). However, we are not aware of published work on aphids that shows impacts of endosymbiont transinfections on plant virus transmission.

So where to next? The BYDV virus inhibition work can be expanded in several ways that can lead to an understanding of both the scope of the transmission inhibition and the mechanisms that might be involved. We are particularly interested in considering the tissue distribution of the *Regiella* and BYDV to see if virus load is reduced in salivary glands from infected aphids. Transcriptomic and genomic analyses could be undertaken to test how the infection might influence host gene expression that contributes to both fitness benefits and virus blocking. Potential impacts on viral transmission under field conditions are also needed, along with tests of specific associations between viral density in plants and acquisition/transmission potential. Beyond this work it would be worthwhile to consider the impact of other *Regiella* isolates on *R. padi* fitness, to consider whether the results described here extend to other clones of *R. padi*, and to consider conditions under which vertical transmission of *Regiella* is stable within lines.

## Acknowledgments

We thank Monica Stelmach and Kelly Richardson for technical assistance and the Grains Innovative Park for supplying the BYDV strain. This work was undertaken as part of the Australian Grains and Horticulture Pest Innovation Program (AGHPIP), supported through funding provided mainly by the Grains Research and Development Corporation (UOM1906-002RTX; UOM2404-006RT), and also by Hort Innovation Australia (ST23002) and the University of Melbourne. This work was also supported by the Scientific and Technological Achievement Transformation Project of the Sichuan Academy of Agricultural Sciences (2024ZSSFGH04).

## Supplementary Material

**Fig. S1.**
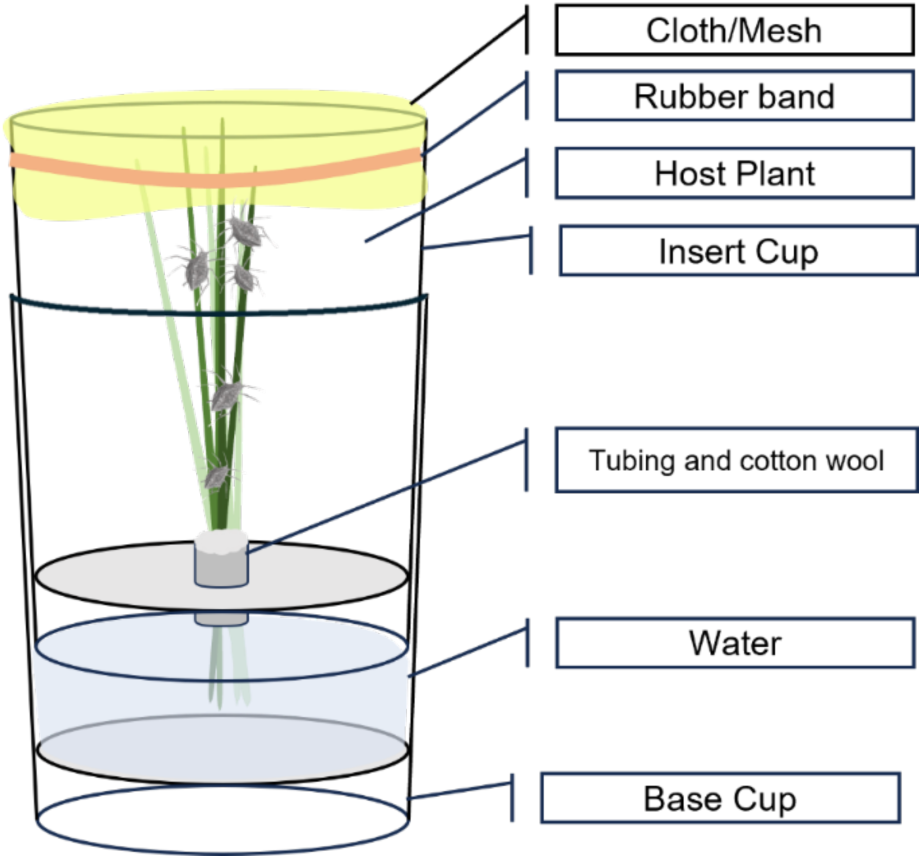
Aphid rearing method using double cups modified from Singh et al. (2020). Five cut wheat plants (GS11-12) submerged in water to keep the plant material from wilting and trimmed to the cup size. The double cup separates the water from the aphid material preventing drowning and high moisture internally. The insert cup allows ease of removal from the base and manipulation for observation of aphid population.

